# Scalable data analysis in proteomics and metabolomics using BioContainers and workflows engines

**DOI:** 10.1101/604413

**Authors:** Yasset Perez-Riverol, Pablo Moreno

## Abstract

The recent improvements in mass spectrometry instruments and new analytical methods are increasing the intersection between proteomics and big data science. In addition, the bioinformatics analysis is becoming an increasingly complex and convoluted process involving multiple algorithms and tools. A wide variety of methods and software tools have been developed for computational proteomics and metabolomics during recent years, and this trend is likely to continue. However, most of the computational proteomics and metabolomics tools are targeted and design for single desktop application limiting the scalability and reproducibility of the data analysis. In this paper we overview the key steps of metabolomic and proteomics data processing including main tools and software use to perform the data analysis. We discuss the combination of software containers with workflows environments for large scale metabolomics and proteomics analysis. Finally, we introduced to the proteomics and metabolomics communities a new approach for reproducible and large-scale data analysis based on BioContainers and two of the most popular workflows environments: Galaxy and Nextflow.

## Introduction

The progress in the application of mass spectrometry (MS) to biological compounds has revolutionized the field of biology: the large-scale identification of proteins and metabolites provides a unique snapshot of a biological system of interest at a given time point [1]. The MS-based high-throughput technologies have resulted in an exponential growth in the dimensionality and sample size [2]. This increase has two major directions: I) the number of samples processed, powered by new mass spectrometers; and II) the number of molecules (metabolites, peptides, and proteins) identified alongside each sample [3]. In addition, the data analysis in MS-based metabolomics and proteomics is becoming more complex, including several convoluted steps to go from the spectra identification to the final list of relevant molecules. This scenario creates major challenges for software developers and the bioinformatics community: I) software and data analysis scalability; II) software availability and findability; and III) reproducibility of the data analysis [3, 4].

Computational proteomics and metabolomics have been dominated by desktop and monolithic software for the past decades, which hampered high throughput analysis in High-Performance Computing systems (HPCS) and cloud environments [5, 6]. Furthermore, many of these are proprietary closed-source solutions, often run only on MS Windows or from vendor’s hardware, and use proprietary binary formats for data intake. These are barriers for reproducible science. During the last decade, open source software and distributed solutions have slowly made their way in these computational fields, with an ecology of computation tools flourishing in proteomics and metabolomics (see reviews in both fields [6, 7]). While open source and distributed frameworks irruption into the aforementioned omics fields is positive for the scalability, portability, and reproducibility of data analysis in this fields, it often comes at a cost of an increased technical complexity: installing, maintaining and executing these analysis software is usually complex and requires advanced software expertise, which is often a rare skill among scientific practitioners. This is further complicated by the fact that reproducibility and collaboration demand the installation of these tools on different computational environments (local computers, HPCS, cloud, collaborators cluster, etc), often requiring different installation processes and software dependencies to be fulfilled [8].

In the past few years, the use of software containers and software packaging systems has markedly increased in general in the field of Bioinformatics [8, 9]. In particular, the BioContainers [8] (http://biocontainers.pro) and BioConda [10] (http://bioconda.github.io) communities have widely increased the availability of containers and adequately packaged bioinformatics tools respectively, providing today thousands of tools in a format that can be used in local workstations, HPCS and cloud environment seamlessly [9]. These software containers reduce the technical entry barrier for setting up scientific open source software and for making setups portable across multiple environments.

While containers and software packages make easier the installation and portability of bioinformatics tools, they still leave to the scientist the task of dealing with combining (plumbing) tools together to create bioinformatics data analysis workflows and pipelines [11]. This is a complex task and demands the use of the Linux command line environment; the underlying file system and data streams. In addition, if the analysis is aimed to run in distributed architectures (e.g. HPC clusters or Cloud), the bioinformatician will need to combine the workflow design (what tools to run with which data inputs and parameters) with the execution logic (e.g. job scheduler, data filesystem). To facilitate workflow design and their execution on different distributed execution environments, such as HPC or Cloud architectures, the bioinformatics community has developed various Bioinformatics workflows systems [11]. During the past 10 years, open source workflow environments have started to consolidate in the field of bioinformatics, and more recently in the past 5 years, these have made an entrance into metabolomics and proteomics. The first popular workflow environment systems in bioinformatics where Taverna (now Apache Taverna) and Galaxy [12], released in 2003 and 2005 respectively. The list is increasing every year with prominent examples such as Nextflow [13], Cromwell, toil and Snakemake [14], among others. Besides workflow systems, proposals for workflow lingua francas have appeared, such as CWL or WDL, among others.

In this manuscript, we will discuss the combination of software containers with workflows environments for large scale metabolomics and proteomics analysis. The combination of software containers and workflows environments promises to make scientific analysis pipelines scalable, reproducible, portable and accessible to scientists that do not have any expertise in the use of complex computational infrastructure and command line environments. We will introduce to the proteomics and metabolomics communities a complete ecosystem of tools and framework for reproducible and large-scale data analysis based on BioContainers and two of the most popular workflows environments: Galaxy [12] and Nextflow [13].

### Current approaches for computational mass spectrometry

In proteomics, the most common strategy for the interpretation of data-dependent acquisition (DDA) MS/MS spectra consists of comparing the experimental spectra to a set of ideal spectra (also called theoretical spectra) extrapolated from the predicted fragmentation of peptides derived from a protein database [15]. During this process, every spectrum obtained by the mass spectrometer needs to be compared with all the theoretical spectra within the same precursor mass. As more data is generated (larger cohorts and more complex samples) the running time becomes longer [16]. During recent years, algorithms and tools have been developed to perform the identification step, such as Andromeda [17], MSGF+ [18] or MSFragger [19]. Even though most of these algorithms have become robust and reliable, analysis of large scale experiments will still be computationally intensive and take considerable execution time [20]. After the identification process, the resulting peptide-spectrum matches can be reliably controlled by false discovery rates filters (such as FDR) (**Figure 1**). Recently, many tools have implemented a secondary database search which takes the initial identification results and refines the search parameters. Finally, the list of quantified protein is ensemble based on the identified peptides by using protein inference algorithms [21, 22]. The list of quantified proteins is provided to the downstream statistical analysis step, which reports the final relevant proteins.

**Figure 1:**
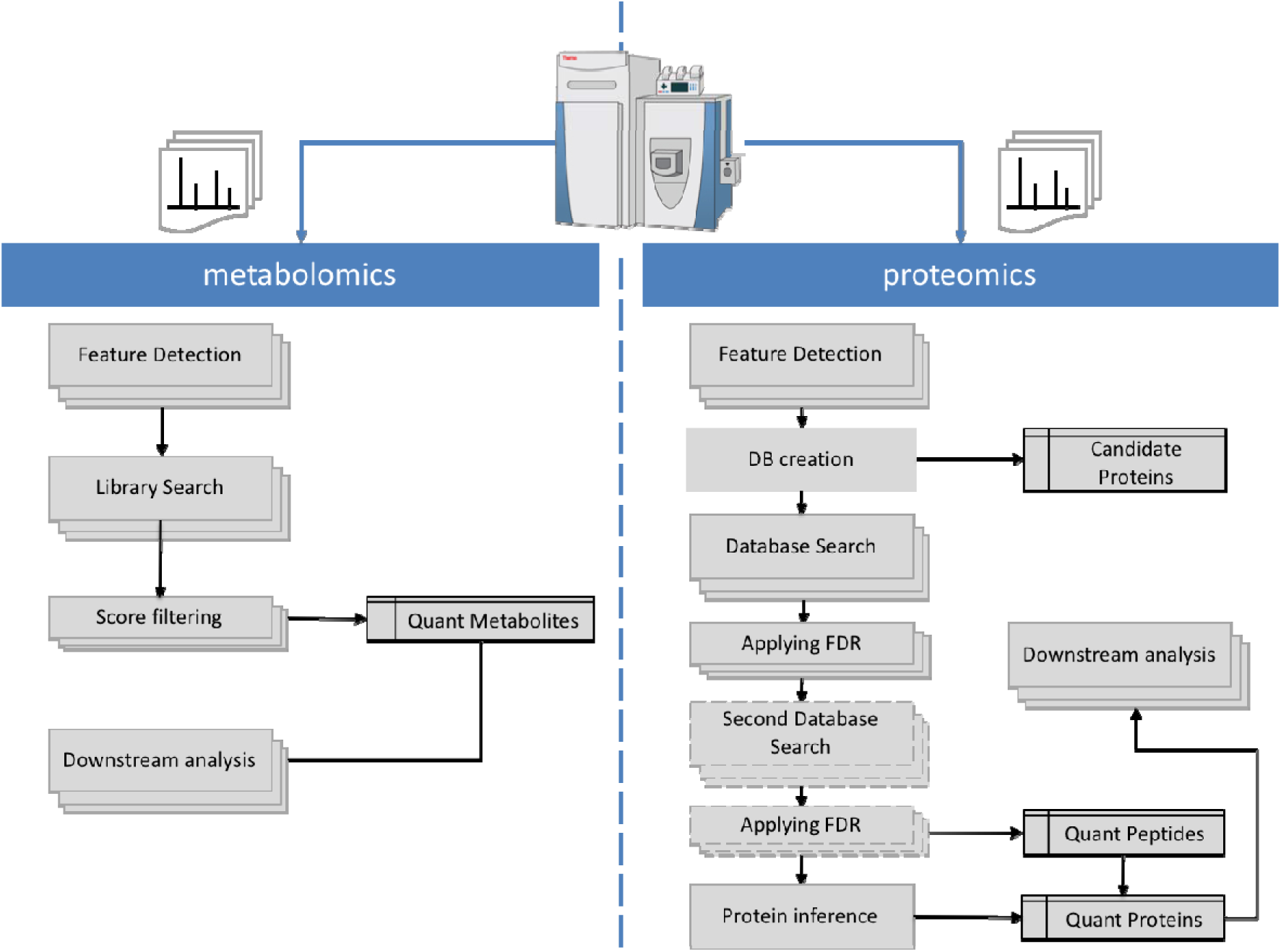
Mass spectrometry metabolomics and proteomics bioinformatics workflow. The Proteomics lane (right) represent a Database search Label-free analysis workflow including Feature detection on MS1 spectra, protein database creation, database search, statistical analysis and final protein inference step. The metabolomics workflow represents a common spectral search workflow.

Data independent acquisition (DIA) is a relatively new mass spectrometry-based technique for systematically collecting tandem mass spectrometry data. Whereas data-dependent acquisition (DDA) selects precursor ions according to their abundances, DIA aims to implement a parallel fragmentation of all precursor ions, regardless of their intensity or other characteristics, enabling the establishment of a complete record of the sample [23]. This analytical method is well-suited for applications requiring the measurement of thousands of proteins or demanding the flexibility to investigate multiple hypotheses without having to acquire additional data sets. Different software have been implemented to analyze DIA datasets such as OpenSWATH [24] and Skyline [25].

Similarly, computational metabolomics is mainly based on the comparison of the metabolites spectra against a well-curated database of previously identified metabolites (Spectral library strategy) (**Figure 1**). Spectral libraries such as METLIN and MassBank contain information about mass and structure of small molecules, although MS/MS spectra are available for only a share of the small molecules in the database. The basic analytical workflow yields thousands of molecular features within minutes of data acquisition. But, similarly to proteomics, only a minority of detected masses can be matched to a molecule in the database, or more commonly to several possible molecular formulas [26, 27]. A statistical validation and manual curation can only be achieved by a matched MS/MS spectrum and/or by another compound-specific property such as retention time, which is then compared to a synthesised standard compound. In principle, quantitative analysis in metabolomic experiments is very similar to the label-free quantitation approaches based on extracted ion chromatograms in proteomic workflows. Feature alignment and detection is followed by quantitation and then perhaps identification of a compound. However, the tendency of small organic molecules to form multimers or adducts (i.e. sodium or ammonium) needs to be considered and detected masses and their intensities deconvoluted before quantitation and statistical evaluation [26].

### Current software ecosystem for computational mass spectrometry

The more established and common tool design for proteomics and metabolomics data analysis are monolithic desktop applications. In this type of bioinformatic tools, all the analysis steps (**Figure 1**) are encapsulated into the same application, which lends itself to be used as a black box, with little understanding from users on the intermediate analysis steps.

### MaxQuant (Proteomics)

MaxQuant [28] is one of the most frequently used platforms for mass-spectrometry (MS)-based proteomics data analysis. The platform includes a Database search engine (Andromeda) to perform the peptide identification and a set of algorithms and tools designed for quantitative label-free proteomics, MS1-level labelling, and isobaric labelling techniques. Recently, MaxQuant has implemented the full export to mzTab file format, enabling the proteomics community to perform complete submissions to ProteomeXchange repositories and analyse the data in an standard file format [29].

In 2013, the MaxQuant team published a detailed documentation of the running time and input/output operations for each step of the analysis [30]. The results showed bottlenecks in overall performance and time-consuming algorithms related to peptide features detection in the MS-1 data as well as the fragment spectrum search. The MaxQuant algorithms are efficiently parallelized on multiple processors and scale well from desktop computers to servers with many cores. However, all the framework and algorithms have been designed as a monolithic tool where all the steps of the data processing cannot be easily distributed in HPC or Cloud architectures. In addition, MaxQuant is not an open source software, which limits the possibility of external contributors and collaborators to reuse the individual components and algorithms moving towards a distributed architecture.

### Skyline (Proteomics)

The Skyline [25] is an open source platform for targeted and data-independent proteomics and metabolomics data analysis. It runs on Microsoft Windows and supports the raw data formats from multiple mass spectrometric vendors. It contains a graphical user interface to display chromatographic data for individual peptide or small molecule analytes. Skyline supports multiple workflows, including selected reaction monitoring (SRM) / multiple reaction monitoring (MRM), parallel reaction monitoring (PRM), data-independent acquisition (DIA/SWATH) and targeted data-dependent acquisition. Because both SRM and DIA data are based on the analysis of MS/MS chromatograms (selected and extracted respectively), the processing (chromatogram peak integration) and visualization of data acquired using these two methods very similar within Skyline. In a recent publication, the Skyline team has recognized that one of the areas to work in the future is the parallelization and distribution of computation and processing in HPC and cloud architectures [31]. These developments will be vital in obtaining the robust, sensitive quantitative measurements required to better understand the systems biology of cells, organisms, and disease states.

### XCMS2 and MZmine2 (Metabolomics)

XCMS-2 [32] and MZmine2 [33] have become arguably the most widely used free software tools for pre-processing untargeted metabolomics data. The XCMS-2 software is publicly available software that can be used within the R statistics language [32]. XCMS-2 is capable of providing structural information for unknown metabolites. This “similarity search” algorithm has been developed to detect possible structural motifs in the unknown metabolite which may produce characteristic fragment ions and neutral losses to related reference compounds contained in reference databases, even if the precursor masses are not the same. In addition, XCMS provides algorithms and tools to find peaks, align/group peaks, correct retention times between different samples, fill peaks, filter by dilution, among other methods [32].

MZmine was first introduced in 2005 as an open-source software toolbox for LC-MS data processing. The first version of MZmine defined the data analysis workflow and implemented simple methods for data processing (e.g. peak noise detection) and visualization. In 2010 [33], a critical assessment of the tool detected that MZmine was a build in a monolithic design, thus limiting the possibility of expanding the software with new methods developed by the scientific community. In lieu of this, MZmine2 was completely redesigned to be modular. MZmine2 was built in multiple data processing modules, with emphasis on easy usability and support for high-resolution spectra processing. MZmine2 includes the identification of peaks using online databases, MS-n data support, improved isotope pattern support, scatter plot visualization, and method for peak list alignment based on the random sample consensus (RANSAC) algorithm.

In 2017, Weber and co-workers conducted a survey on software data usage in metabolomics and found that LC-MS data analysis in metabolomics is performed in 84% of the cases using open-source tools. The predominant open-source software is XCMS (70%), followed by Mzmine and MZmine2 (26%). Interestingly, most of the usages of XCMS is through the Online XCMS Portal (https://xcmsonline.scripps.edu/landing_page.php?pgcontent=mainPage) a popular Web application that helps the user to go through each step of the data analysis. XCMS Online directly searches the experimental mass spectra into METLIN online data using the traditional precursor ion selection window and additionally a distance matrix score to obtain good spectral matches. For the reader interested in a wider landscape of Metabolomics tools and their usage in different scenarios, Spicer et al^6^ provides guidance.

### From desktop applications to distributed HPC and cloud architectures

The field of MS-based computational proteomics and metabolomics heavily relies on monolithic desktop applications which prevents the analysis of the big amount of data in HPC clusters and cloud architectures. In order to overcome these problems, three main fields of computer science and algorithm development need to develop and grow in the near future in computational MS-based proteomics and metabolomics: i) component-based and modularized applications; ii) standard practices for tool packaging and deployment, and iii) workflow systems.

**Figure 2** shows a proposal for the next-generation of computational proteomics tools and algorithms. Each step of the data processing should be designed as an independent module component that can be executed in an independent node distributed architecture. Multiple tools and modules can be packaged into the same framework (e.g. OpenMS [34]) but all of them should be executable independently and interchangeably (through adequate exchange intermediate formats). A component-based software framework is a branch of software development that emphasizes the separation of concerns with respect to the wide-ranging functionality available throughout a given software system. The component-based development allows the data analyst to replace and substitute components of the workflow by new tools or a new version of the same tool without impacting the development process.

**Figure 2:**
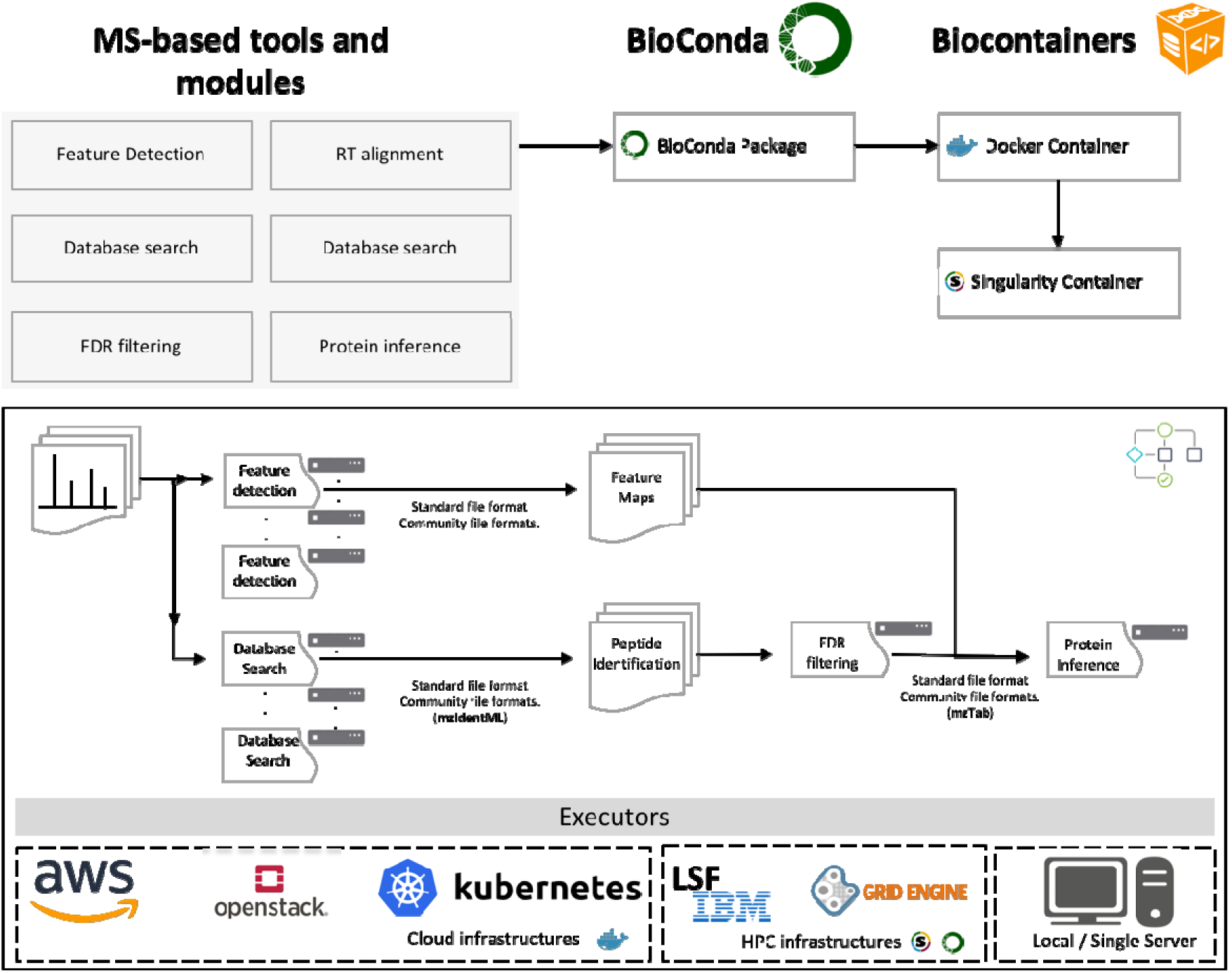
The proposed roadmap to scale metabolomics and proteomics data analysis includes the packaging and containerization of the specific tool and software using BioConda and BioContainers. The design of bioinformatics workflows that use the specific containers and abstract the execution from the compute environment (e.g. Cloud or HPC). A very important step of this design is the use of standard file formats that enable to communicate different tools and steps of the workflow.

In MS-based proteomics three different frameworks have been designed from the beginning on component-based architecture: OpenMS [34], Trans-proteomics pipeline [35] and OpenSWATH [24]. OpenMS and OpenSWATH provide a set of computational tools which can be easily combined into analysis pipelines even by non-experts and can be used in proteomics workflows. These applications range from useful utilities (file format conversions, peak picking) to wrappers for known applications like peptide identification search engines. These two frameworks have been used recently to analyse big datasets [36, 37]. Though these frameworks have been fully implemented as component-based frameworks, they have been really slow to implement and promote standard file formats between each component.

### Standard file formats for better compatibility between components

Standard file formats allow developing a common persistence (e.g. file) representation of the data that is analysed (e.g. spectra, peptides). The proposed approach in **Figure 2** reduces the need for components to translate from one file to another, which increase input-output (IO) operations and data transformation. This standard file format enables the development of new tools based on a common representation making the input and output parameters of tools common to any software component.

The Human Proteome Organization (HUPO) and the Proteomics Standards Initiative (PSI) has developed for 15 years file formats and common representations for MS-based proteomics and metabolomics data, from the spectra to protein expression [38]. For mass spectra, the mzML specification is the most stable, robust and mature file format, representing not only the MS/MS signal but also chromatograms and instrument metadata [39]. For peptide/protein identification results, the mzIdentML file format not only captures the peptide and protein identifications but also the software metadata (e.g. FDR thresholds, search parameters) use to perform the analysis [40]. Finally, mzTab file format store the information of quantitative data for proteomics and metabolomics experiments [41, 42].

Despite advances in the past years, standard file formats need time to evolve and consolidate. The software development process shouldn’t be slower because of the development of a specific file format. Our recommendation is to replace and reuse existing file formats when is possible and avoid the creation of new ones. A good example of these efforts is the peptide search engine MSGF+ that natively use mzIdentML and has been extensively used in open source workflows.

A major problematic in standardised workflows for metabolomics and proteomics in the field of mass spectrometry is the lack of intermediate exchange formats similar to the existing genomics formats (such as BAM, SAM, CRAM, VCF, bed, etc. to name a few). Often in these younger fields tools will generate results in ad-hoc formatted files with poor specifications and often incompatible with downstream tools that would naturally pipe. This means that further tailored conversion steps need to be provisioned, which slow development, require maintenance, and might in cases introduce errors or data loss.

### Packaging and deployment using BioContainers

A component-based architecture like the one proposed in this manuscript (**Figure 2**) prompts multiple challenges in deployment and execution. Moving from single desktop applications to distributing data analysis and complex workflow systems creates major challenges for the bioinformatics community: (i) software availability, (ii) results reproducibility and (iii) automated software/environment deployment. Component-based pipelines require substantial effort for correct installation and configuration (e.g. conflicting dependencies, different base libraries). In addition, versioning components, key for reproducing the results of the analysis, is a burden to scientific software development groups, who are less used to proper software engineering standards.

Recently, containers and packaging technologies such as Conda (http://conda.io), Docker (http://www.docker.com) or Singularity (https://www.sylabs.io/) have emerged to overcome these challenges by automating the deployment of applications inside so-called software containers. The BioContainers community [8] (http://biocontainers.pro) has created a complete architecture and solution to overcome these challenges based on community-driven BioConda packages [10] and Docker containers.

The BioContainers community has defined a set of guidelines about how to create containers, deploy and maintain them (**Figure 3**) [9]. Each component (software tool) developer can create a Conda recipe (*a set of yaml and bash scripts which describe how to consistently install a software package on Linux*) or a Docker build recipe (*Dockerfile*), which are all stored in GitHub. Each contribution (new recipe) is accepted using a Pull Request (PR) mechanism by GitHub service [43]. These Pull Requests are accepted by members of the bioinformatics community that get given permissions, reducing the burden on a small group of maintainers and hence making the model sustainable. After the PR get merged and the recipe merges into the repository with a well-defined version, a continuous-integration system is triggered, creating the Conda package (in case of a Conda recipe) and the corresponding Docker container and Singularity containers tagged at that version. Historic versions of the same package are stored both as Conda packages and containers, guaranteeing future reproducibility of older pipelines that use earlier versions of a tool.

**Figure 3:**
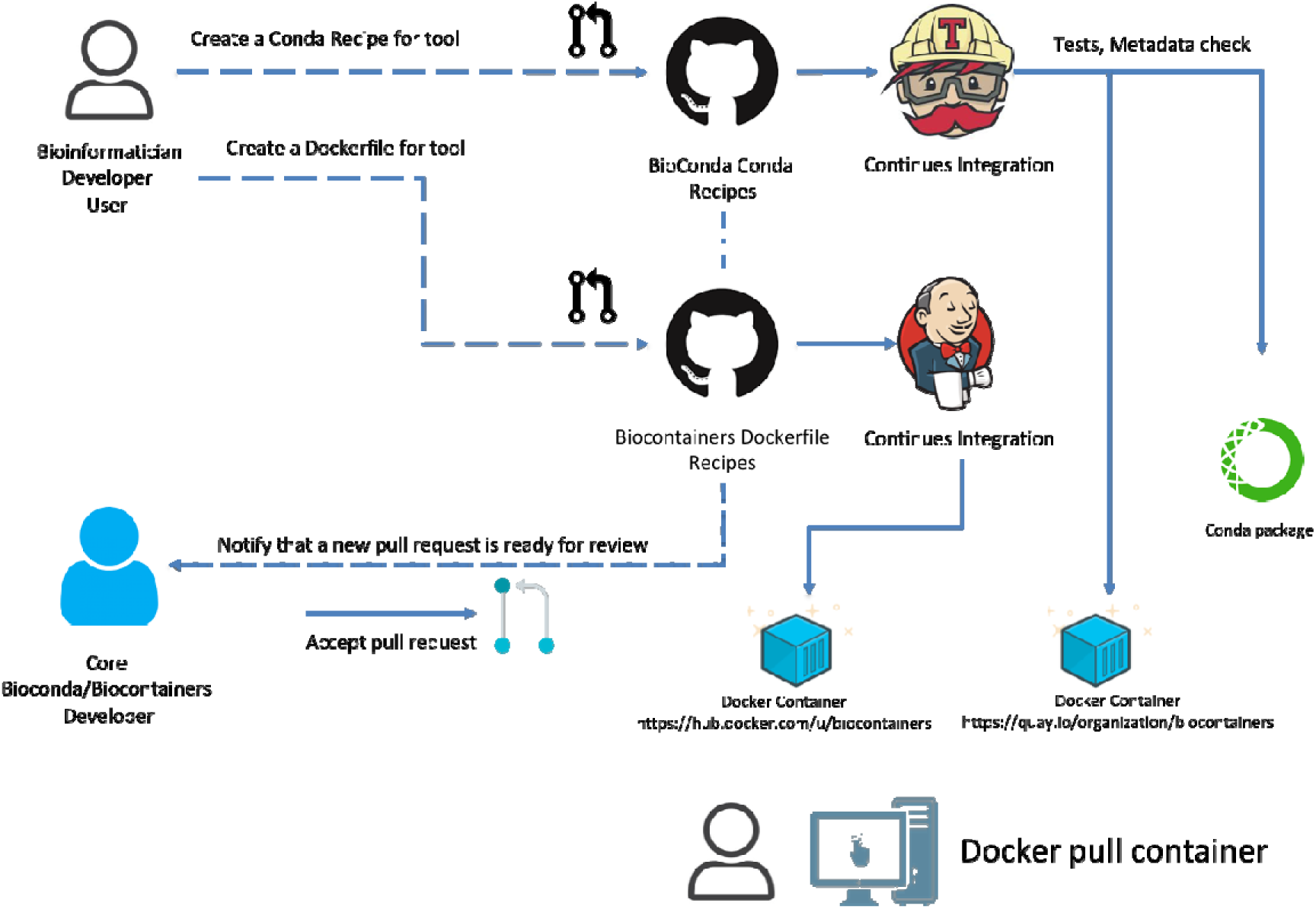
BioContainers architecture from the container request by the user in GitHub to the final container deposited in DockerHub (https://hub.docker.com/u/biocontainers) and Quay.io (https://quay.io/organization/biocontainers). The BioContainers community in collaboration with the BioConda community defines a set of guidelines and protocols to create a Conda and Docker container including: mandatory metadata, tests, trusted images [9]. The proposed architecture uses a continues integration systems (CI) to test and build the final containers and deposited them into public registries. All the Containers and tools can be searched from the BioContainers registry (http://biocontainers.pro/regitry).

The created package and containers contain all software dependencies needed to execute the tool in question. In general, one package will contain simply one tool, large packages containing many tools are in general discouraged. This allows to execute the pipeline in different compute environments, without the complexity of installation, dependency management, etc. It also allows moving the pipeline from one environment to another (e.g. HPC, Cloud or local personal computer) because everything is executed in containers. At the time of writing, BioContainers provides more than 7000 bioinformatics containers that can be searched, tagged and accessed through a common web registry (https://biocontainers.pro/#/registry/). Importantly, the BioContainers and BioConda communities convert automatically Bioconductor packages automatically into containers.

### Workflow systems

High-throughput bioinformatic genomics and transcriptomics analyses increasingly rely on pipeline frameworks to process sequence and metadata. Until recently, this was not the case for Proteomics and metabolomics, where mostly the analysis happens on single desktop machines. Modern workflows systems such as Galaxy [12], Cromwell (https://github.com/broadinstitute/cromwell), CWL tool (https://www.commonwl.org/v1.0/CommandLineTool.html), Toil [44], Nextflow [13] or Snakemake [14] frameworks are playing an important role to move the data analysis steps from the single desktop applications into distributed compute platforms [11]. All these workflow engines provide three four major functionalities for data processing: i) execution in distributed architectures (HPC, Cloud); ii) separation between thee execution environment and workflow design; iii) recovery/restart mechanisms for failed components and tasks; iv) support for automatic tool installation using Conda, Docker or Singularity technologies. This last feature (automatic tool resolution using packaging systems) allows developers and bioinformaticians to execute the workflows and pipelines without the need to install and configure each tool manually in the desired architecture. Two different workflows system with a lot of attention from the Bioinformatics community are NextFlow [13] and Galaxy [12].

### NextFlow

NextFlow (https://www.nextflow.io/), an expressive, versatile and particularly comprehensive framework for composing and executing workflows. NextFlow uses a domain-specific language (DSL) which also supports the full syntax and semantics of Groovy, a dynamic language that runs on the Java platform. One of the great features that make NextFlow a powerful workflow engine is its dataflow functionalities. Nextflow allows users within the workflow definition to filter data, run processes conditionally on data value or have splitting/merging pipeline steps expressed in a short, elegant syntax.

Nextflow separates the workflow definition from the execution environment, which allows users to execute the same workflow in different architectures (Cloud, HPC or a local machine). This abstraction level is gurranted by using an execution layout that defines which type of containers will be used to execute the tools (components of the workflow) and which type of architecture will be used to execute those containers (e.g. HPC, Cloud). Currently, Nextflow supports the following technologies: Conda, Docker and Singularity; and the following execution environments: Local (the software run in single node, server), HPC clusters including (Sun Grid Engine, IBM LSF, PBS/Torque, HTCondor) and cloud providers, such as Amazon Web services (through AWS batch) or Google Cloud Platform (see full list here: https://www.nextflow.io/docs/latest/executor.html). This execution layout can be configured with workflow variables which enable to switch between architectures with no hassle.

**Figure 4** shows an example of Nextflow workflow with one process to perform sequence alignment using Blast (container https://biocontainers.pro/#/tools/blast). Nextflow allows bioinformaticians to perform analysis in different architectures with the same workflow definition (https://www.nextflow.io/docs/latest/basic.html). Each step of the workflow (called process) describes which process will be performed and the input/output parameters. The container section inside the process (*blastSearch*) states which containers will be used; including container name (*blast*), version of the container (*v2.2.31_cv2*). Between triple quotes is the actual command will be executed in the container (in this case blast). This is needed because one container can provide multiple tools. Finally, the Nextflow config file (https://www.nextflow.io/docs/latest/config.html) defines how the present workflow will be executed. In the example, we have defined two possible scenarios: *local* and *lsf*. If the user runs the workflow using the local configuration (command - *nextflow workflow.nf -c config.nf -profile local*) it will be using BioContainers Docker containers, if the user uses *lsf* (command - *nextflow workflow.nf -c config.nf -profile lsf*), then will be using BioContainers singularity and the LSF cluster executor.

**Figure 4:**
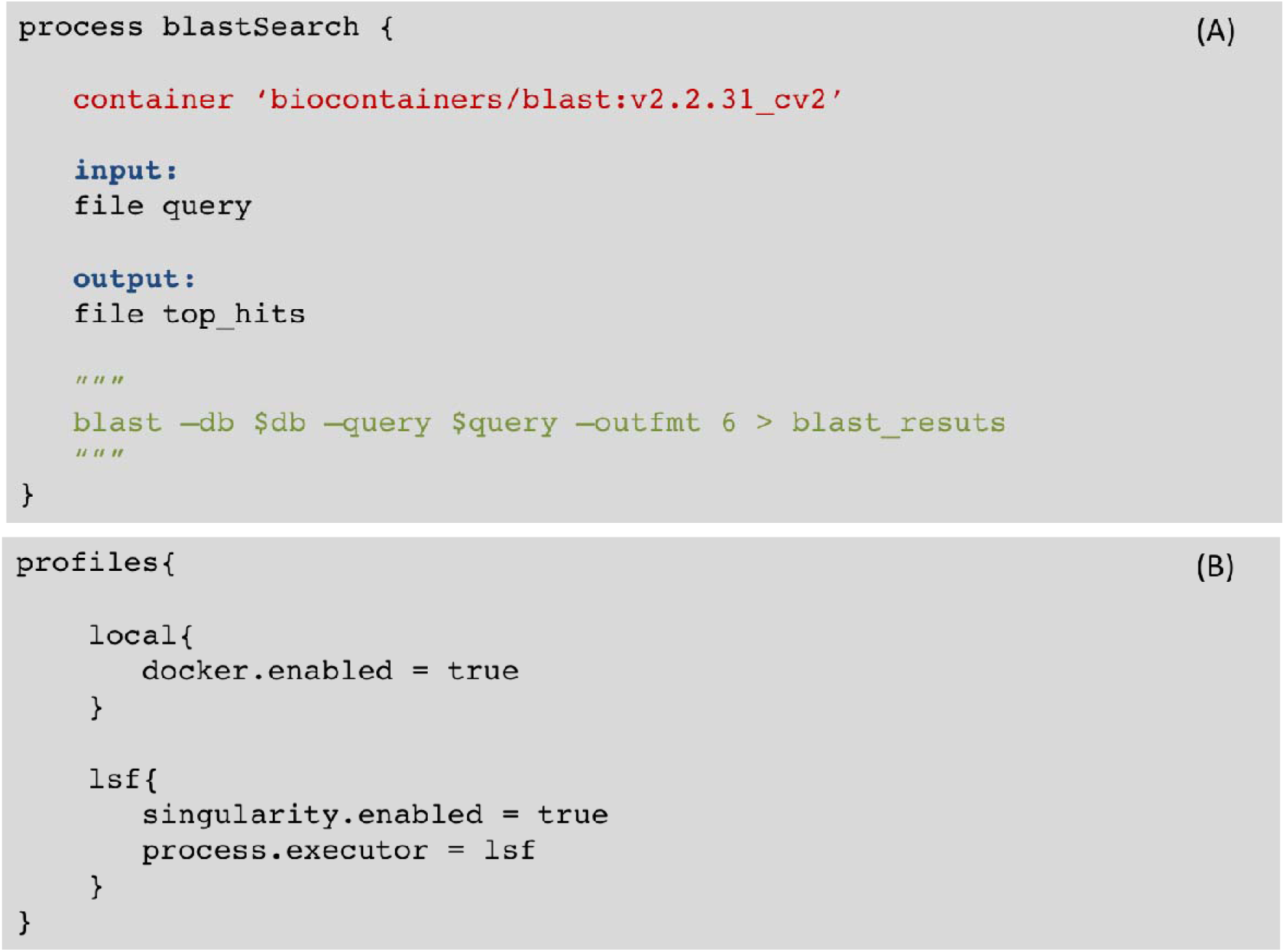
Nextflow allows bioinformaticians to perform analysis in different architectures with the same workflow definition. **(A)** The workflow step (called process) describes which process will be performed and the input/output parameters. The *container* section inside the *blastSearch* process state which containers will be use; including **container name** (blast), and **version of the container** (v2.2.31_cv2). Between triple quotes is the actual command will be executed in the container (in this case blast). This is needed because one container can provide multiple tools. **(B)** The Nextflow config file (https://www.nextflow.io/docs/latest/config.html) defines how the present workflow (A) will be executed. In the example, we have defined two possible scenarios: local and *lsf*. If the user run the workflow using the local configuration it will be using Docker containers, if the user uses *lsf*, then will be using singularity and the LSF cluster executor.

A list of Nextflow and proteomics workflows using BioContainers can be found in this GitHub repository (https://github.com/bigbio/nf-workflows/). In addition, it provides configuration variables to customize the computer/hardware that is required to perform each task. For example, the user can customize the type of node (Memory, number of cores) that are needed for each specific task (component tool). Another important feature of NextFlow is the simplicity of the language syntax and the support of workflow versioning, which enables better reproducibility.

### Galaxy Project

Galaxy (https://galaxyproject.org/) is a web application workflow environment written in Python, capable of distributing jobs among a plethora of batch schedulers (PBS, LSF, GoDocker, DRMAA based schedulers, etc), local machine and cloud providers (through Kubernetes [45] and others). In the HPC cases, Galaxy provides the flexibility to use either containers (Docker or Singularity) or directly Conda packages. Galaxy tool wrappers are written to point to specific package versions, towards reproducibility. Galaxy provides a complete separation of concerns between the workflow logic definition and the actual execution. It offers reusability as well at the tool level, meaning that not only workflows can be shared but also individual tools. Both tools and workflows are versioned in Galaxy (and multiple versions of a tool can be installed on the same instance). Galaxy provides tool/workflow repositories, called Toolsheds, where users can deposit and find currently more than 6,000 tools wrapped for Galaxy, and automatically install desired versions of those tools to their own Galaxy instance. Tool’s dependencies are resolved automatically by Galaxy using either Conda packages, Docker or singularity containers, depending on setup. On the same workflow, different tools can be sent to different underlying executors and rely on different dependency resolution as well. Besides a rich and responsive user interface (UI), Galaxy allows operations through a mature REST API, Python clients (e.g. bioblend, ephemeris) and command line interface (parsec), to programmatically control the execution of tools/workflows and data upload/downloads. Galaxy fulfils the promise of a single workflow environment system for research, training, and production environments. The same system can satisfy the need of researchers with no bioinformatics expertise – but in need of doing data analysis through a UI – or tenured bioinformaticians wishing to systematize the execution of production pipelines through a CLI.

Many organizations provide computing power to end users in the need of doing biological data analysis through public Galaxy instances -- in the region of 100 public instances exist today -- which are normally flavoured for different research topics. Notable instances in terms of size are usegalaxy.org (http://usegalaxy.org) and usegalaxy.eu (http://usegalasy.eu). Galaxy is organized in different initiatives that help to download and deploy complete solutions of galaxy tools for a specific field (e.g. Proteomics).

The Galaxy-P (Galaxy for Proteomics - http://galaxyp.org/) initiative provides workflows and tools in the fields of proteogenomics and metaproteomics (http://galaxyp.org/access-galaxy-p/). PhenoMeNal [46] (https://public.phenomenal-h2020.eu/), Galaxy-M [47], and Workflow4Metabolomics [48] (https://workflow4metabolomics.org/) are the most complete compendium of tools and workflows available in Galaxy for metabolomics researchers.

### Towards reproducible data analysis

Reproducibility is challenged in life sciences, especially in computationally intensive domains (e.g. proteomics and metabolomics) where results rely on a series of complex analytical and bioinformatics steps that are not well captured, by traditional publication approaches. While there are now several guidelines and platforms to enable reproducibility in computational biology [43, 49, 50], the approach we describe here is flexible, robust and scalable enough to guaranty the features for reproducibility research: (i) managing software dependencies, (ii) separation between the data analysis design and the local computational environments, and (3) virtualizing entire analyses for complete portability and preservation against time [49].

The use of BioConda and BioContainers as independent components in data analysis resolves the problem of complex software dependencies. In addition, it provides a mechanism to easily replace independent components from different technologies and programming languages (e.g. python by R package). The use of workflows in combination with container technologies allows researchers to reproduce data analysis in their own compute architecture (e.g. local PC or cloud). BioConda and BioContainers provides a consistent versioning system and combined with virtualization allows to complete entire data analysis overtime.

Finally, in order to complement the software efforts made by the BioConda and BioContainers communities, we urge software developers of the metabolomics and proteomics communities to embrace standard file formats as supported input and output of each software and component tool. Standard file formats not only enable better interoperability between software components, but also an improve the reproducibility of the analysis [51].

## Conclusions

Proteomics and metabolomics mass spectrometry are moving from desktop application data analysis to distributed architectures (HPC and Cloud) due to larger datasets being generated (more sample, more replicates, higher coverage, more resolution, etc). However, the major software used in the field, such as Skyline, MaxQuant, ProteomeDiscover and XCMS Online, are mainly developed as monolithic tools, hampering the scale up of individual steps of the analysis into distributed architectures. First, the software development and algorithm paradigm should be changed by decoupling monolithic applications into smaller component (tasks) tools that can be distributed on Cloud and HPC architectures. We recommend that each of these small components support standard file formats for inputs and outputs, towards facilitating the exchange of steps in data analysis pipelines.

We presented a future paradigm for proteomics and metabolomics large scale data analysis based on BioContainers, BioConda, Docker/Singularity containers and workflow engines such as Galaxy and Nextflow. The proposed idea starts by creating a Conda or Dockerfile recipe in BioContainers providing the mandatory metadata and dependencies to build the container. The new recipe will be built using continuous integration where the BioContainers architecture will check the metadata, tests and push the final containers into BioContainers public registries. The Conda-based containers are deployed in Quay.io (https://quay.io/organization/biocontainers) and the Dockerfile-based are deployed in DockerHub (https://hub.docker.com/u/biocontainers). All containers are searchable, and discoverable through the BioContainers tool registry (http://biocontainers.pro/#/registry/).

Finally, we recommend the proteomics and metabolomics community to embrace the development of bioinformatics workflows and gradually move bioinformatics pipelines and data analysis into workflow environments such as Nextflow or Galaxy. The combination of workflow environments and BioContainers will enable more reproducible and scalable metabolomics and proteomics data analysis.

## Acknowledgments

Y.P.R is supported by BBSRC Grant “ProteoGenomics” [BB/L024225/1]. We would like to thanks to ELIXIR and the ELIXIR Tools and Compute platforms for the support to BioContainers Community and Architecture BioContainers

## Abbreviations

AWS: Amazon web services
DDA: Data-dependant acquisition
DIA: Data-independent acquisition
DSL: Domain-specific language
FDR: False discovery rate
HPC: High-performance computer
HPCS: High-performance computing systems
HUPO: Human Proteome Organization
IO: Input-output
MRM: Multiple reaction monitoring
MS: Mass spectrometry
MS/MS: Tandem mass spectrometry
LC-MS: Liquid chromatography–mass spectrometry
LSF: IBM Platform LSF
PR: Pull request
PRM: Parallel reaction monitoring
PSI: Proteomics Standards Initiative
REST API: Representational State Transfer programming interface
SRM: Selected reaction monitoring

